# Decreasing alpha flow releases task-specific processing paths

**DOI:** 10.1101/2023.06.18.545474

**Authors:** Jevri Hanna, Cora Kim, Stefan Rampp, Michael Buchfelder, Nadia Müller-Voggel

## Abstract

Directing and maintaining attention toward relevant information and away from non-relevant information is a critical competence of higher-order nervous systems. Here, we used directed connectivity to investigate how the human brain prioritizes appropriate processing paths when participants are performing a behavioral task requiring attention in the visual or the auditory modality. We show that the efficient transfer of information relevant for the task is mediated by a region- and task-specific decrease of alpha band connectivity from parietal and, in case a motor response is required, motor cortex to the relevant sensory cortices. Further, alpha-band connectivity modulations reliably predict alpha power modulations in the task-related sensory cortices, particularly where the task-irrelevant cortex is inhibited via local alpha power increases. We suggest that the task- and region-specific modulation of alpha-band connectivity in the present study is a basic neuronal mechanism orchestrating the allocation of task-relevant neuronal resources related to and possibly underlying the previously reported sensory alpha power modulations in association with the allocation of competing attentional resources.

## Introduction

The adaptive allocation of processing resources and the dynamic integration of information processed in different brain regions as required by a given task or environment is a key facet of human cognition. These attentional dynamics can change by means of bottom-up mechanisms, like stimulus salience in a given sensory stream, but also through top-down influences, such as selective attention for a specific task or goal [1], which is the focus of our present study. We use here oscillatory phase-based, directed connectivity measures on source localized MEG data to observe how attentional allocation is reflected in the functional human connectome. In addition to describing oscillatory network properties of the brain and their changes from resting state into a task state, the present study particularly focuses on connectivity changes in the alpha band as a possible mechanism of fast and flexible top-down prioritization of resources.

### Alpha power and attentional resource management

Previous research has primarily focused on observing changes in local alpha power with attentional tasks. Jensen and Mazaheri [2] proposed in their gating-by-inhibition model that the fast and flexible routing of information flow is realized by the modulation of local alpha power in sensory brain regions, inhibiting task-irrelevant and facilitating task-relevant information processing. This theory obtained support from neurophysiological research across different modalities showing that systematic modulations of ongoing alpha oscillations in sensory brain regions facilitate or inhibit visual, somatosensory and auditory processing during attention or memory tasks [3–13]. Most evidence shows that the decrease of alpha power is related to increased excitability in the relevant brain region and a facilitation of information processing, while the increase of alpha power is associated with a functional inhibition of the accordant brain region resulting in the inhibition of processing and a gating of perception. More recent research suggests that the association between high alpha power in sensory regions and the inhibition of perceptual processing is more complex and critically dependent on the utilized experimental paradigm [14–22]. For instance, alpha power modulations have been observed during anticipation but not during stimulus processing [14] or were completely absent during a perceptual suppression task [18]. Moreover, Zhigalov and Jensen [21] recently showed using the classical visual spatial attention paradigm that alpha power is not necessarily modulated in sensory cortex but rather further down the processing stream in the dorsal parietal cortex. Research has thus revealed a systematic relationship between the level of alpha power in specific brain regions, be it sensory or parietal, and the gating of information flow, albeit not as consistent as initially expected (e.g. [14, 18]).

### Higher order regions influence sensory alpha power

Apart from that, it has not yet been fully understood whether the modulation of alpha power in sensory and parietal brain regions is the fundamental neuronal mechanism determining the allocation of neuronal resources or if the local alpha power modulations are rather an epiphenomenon of more basic processes [15, 23]. Given that alpha power modulations were almost exclusively investigated in single, sensory brain regions, this ambiguity is perhaps not surprising. Conceptualizing attention as the *routing of information flow* in the sense of a dynamic prioritization of relevant processing paths would rather imply neuronal processes acting beyond the borders of specific regions, most likely appearing in specific patterns of connectivity between them.

An essential first step for a more comprehensive understanding of the underlying processes responsible for the allocation of neuronal resources would be to recognize from where and through which channels the local alpha power modulations are initiated [24]. Some previous findings suggest that task-specific higher-order regions might orchestrate the allocation of resources. Kastner and Ungerleider [25] summarize research pointing to higher-order regions or networks generating modulatory influences on sensory cortex with attention. Likewise, it has been shown that the modulation of brain activity in frontal or parietal brain regions induced by Transcranial Magnetic Stimulation (TMS) or lesions in frontal-parietal networks affect alpha power in sensory brain regions, and sensory processing in a task-specific manner [26–29]. It has, however, not been clarified yet how or by which means a higher-order region can prioritize or de-prioritize processing resources of a specific sensory region and influence it to adjust its sensitivity, by e.g. modulating local alpha power.

### Directed connectivity as a possible mechanism of influence from higher-order to sensory regions

It is possible that such top-down influences on processing in sensory regions come into effect by adjusting the routing of information flow in the brain, prioritizing and/or de-prioritizing specific paths in the network by modulating their strength or direction. Several studies suggest that, for the allocation of neuronal resources during attention tasks, communication between parietal and/or frontal and sensory areas are relevant [30–34]. Most interestingly, these studies propose that the transfer of information from higher-order regions to sensory cortices is mediated through the alpha frequency band reminiscent of the local alpha power modulations in sensory cortices themselves. Popov and colleagues [35] showed using Granger causality that parts of the frontal cortex modulated occipital alpha oscillations during a visuospatial attention task. Findings from animal studies corroborate the significance of alpha or beta frequencies for feedback or top-down communication [36, 37]. Also, Lobier and colleagues [31] supported the role of connectivity in the alpha band for the routing of information flow and revealed a synchronization of high-alpha oscillations between frontal, parietal and visual areas during visuospatial attention. They, however, did not investigate the direction of information flow. Wang and colleagues applied Granger causality to high-density EEG data during resting state and tasks requiring attentional selection or memory [33]. The authors report that directed alpha-band connectivity from task-specific right frontal areas to occipital cortex decreased during the visual spatial attention task, while they observed an increase of alpha connectivity from left frontal areas to occipital cortex for the memory task. They conjecture that a reduction of alpha-band Granger causality from higher-order regions to the visual cortex with spatial attention could result in a disinhibition of the visual cortex to increase local excitability. This would be in line with the gating-by-inhibition theory mentioned above [2] and suggest that, for an optimal allocation of resources, task-specific higher-order regions adjust information flow through directed alpha-band connectivity to relevant sensory regions. As these results did not survive correction for multiple comparisons, they should, however, be interpreted with caution. In conclusion, alpha connectivity seems to be relevant for the allocation of resources and the routing of information flow. However, apart from the above-mentioned study by Wang and colleagues [33], a systematic investigation of attentional resource management via directed connectivity from higher-order regions to lower-level sensory regions is still missing, and the question whether connectivity patterns relate to local adjustments of cortical excitability in the form of alpha power modulations is still unanswered.

Studying whole-brain directed connectivity patterns and changes with attentional tasks in a systematic way may allow us to observe more directly how the patterns and changes of information flow contribute to the flexible allocation of resources and to understand more precisely how long-range connectivity dynamics interact with local alpha oscillations in this process. With the present study, we aimed to test if the allocation of resources in the brain is mediated through modulations of directed alpha-band connectivity from task-relevant higher-order regions to task-relevant sensory cortices. To this end, we presented participants with a set of basic sensory attention tasks alternately requiring selective attention to one modality of concurrently present continuous auditory and visual stimuli. Participants had to detect and react to subtle modulations within the attended stream (visual or audio) by a right hand button press. One control condition was a repeated version of the visual attention task with increased difficulty by introducing auditory distractions; a second control task required internal attention away from the external stimuli (while counting backwards) and no motor reaction was required. To assess the routing of information flow as precisely as possible and take into account both its direction and frequency, we applied directed Phase Transfer Entropy (dPTE) to source-localized MEG data. dPTE is a relatively new connectivity measure, which provides a robust estimation of oscillatory information flow from one region to another in a desired frequency band (Lobier et al. 2014) by quantifying how knowledge of the (statistical) distribution of phase at an earlier time point in both a source region and a target region improves prediction of phase distribution in the target region at a later time point, in comparison to the predictive power obtained by using the earlier time point distribution in the target region only. The underlying logic of this approach is very similar to that of Granger Causality, but unlike Granger Causality, dPTE is particularly robust against the noise and linear mixing which is intrinsic to M/EEG data, and makes less assumptions about the relationship between signals [38]. Earlier dPTE research has already revealed characteristic patterns of connectivity during resting state, most prominently from anterior to posterior regions in the theta band (4-8Hz) and from posterior to anterior regions in the alpha (8-13Hz) and beta (13-30Hz) bands [39–42]. In order to gain a complete picture of how the previously described connectivity pattern during resting state [39] changes when entering a task state, we investigated connectivity during rest and tasks for a set of frequency bands (theta (4-8Hz), low alpha (8-10Hz), high alpha (10-13Hz), beta (13-30Hz), and gamma (31-48Hz)). Based on the majority of available literature, however, we expected that top-down attentional processes would modulate primarily the alpha bands. We hypothesized systematic changes in information flow from the resting state pattern when participants perform the attention tasks, and that these would exhibit a task- and region-specific modulation of information flow from higher-order association regions (parietal and/or frontal) to task-relevant sensory regions (visual and auditory). Finally, we planned to test whether systematic changes in connectivity from higher to sensory regions with attention tasks are associated with local alpha power modulations in the sensory cortices, aiming to explore the functional interplay of these two phenomena.

## Methods

### Participants

We obtained MEG data and structural MRIs for 28 normal-hearing participants, recruited from flyers posted online on social media. Three were excluded for excessive head movement during the MEG recording (more than 5mm between blocks), and another participant was excluded for not making any behavioral responses, leaving a total of N=24 (8 male, mean age 25.3, age range 20-37) participants. All participants reported being right-handed, and not under any relevant medication and provided their written informed consent for participation in the study. The procedures of the study were approved by the Ethics Committee of the Friedrich-Alexander-Universität Erlangen-Nürnberg, Department of Medicine (registration number 52_17B).

### Procedure

After receiving instructions, participants were fitted with head position indicator (HPI) coils. Head shape was measured with a digitizer stylus, and participants were then positioned supine in the MEG. Video stimulation was delivered through a mirror and projector system, and audio stimulation was delivered through binaural air tubes terminating in the ear canal. With sound delivery by air tubes, it is usually the case that left and right tubes are unevenly placed in the canal, causing sound to seem louder in one ear than the other. In order to remediate this problem, we administered a hearing test through the tubes, where participants heard the different tones that would later be part of the experiment. Tones were played with varying loudness to the left or right ear and participants answered with a button press of the left or right hand whether they heard a tone on the left or right side. Tone loudness was successively lowered or raised until the hearing threshold was determined for each experimental tone for each ear within a few decibels. The audio stimuli were then each adjusted to 55dB above threshold for the given ear and were thus perceived as equally loud in both ears during the experiment. There was a short practice round where it was confirmed that the participants correctly understood the tasks, and that all equipment was functioning properly. Then the experiment commenced with a 3 minute, eyes-open resting state recording. Afterwards, participants were presented with four different experimental tasks, in counter-balanced order, with a short break in between each one. During these breaks, HPI coils were measured to assess head movement between the experimental blocks. Each of the four experimental tasks lasted about 10 minutes, and participants were generally in the MEG for approximately an hour. The entire procedure, including explanations, preparation, and debriefing lasted around two hours, for which participants were paid 10€ per hour, both for this procedure and for the accompanying structural MRI.

### Experimental tasks

Four experimental tasks were conducted in counterbalanced order. In all tasks, participants perceived constant audio-visual background stimuli consisting of a fixation cross and a continuous sound. All tasks also involved instructions on which modality to focus attention. Three tasks required participants to direct attention to one perceptual modality, and to detect modulations in the stream to which they had to respond with a button press. Of these three tasks, one required attention to modulation detection in the audio domain (audio task), a second one required attention to modulation detection in the visual domain (visual task). A third task required to direct attention and respond to modulations in the visual domain, albeit with constant distractions in the auditory domain that had to be ignored (visual with distraction). This task including distractions was added as a comparison case with increased task difficulty and cross-modal competition. In these three tasks, participants were required to react with a button press of the right hand as quickly as possible after detecting target modulations. In a fourth task, participants were asked to silently count backwards to induce a situation where participants are attending internally and do not react to external visual or auditory stimuli.

All four tasks each consisted of four trials of 100 seconds length of continuous audio-visual stimulation, during which one of four possible sounds was continuously played while a fixation cross was presented. The fixation cross was white, centered on a gray background. Participants were asked to focus their eyes on the fixation cross at all times. The four possible auditory stimuli presented were:

: 1) a 4000Hz sinus tone, 2) a 7000Hz sinus tone, 3) white noise with an FFT gaussian filter centered at 4000Hz, which produced a sound similar to crickets at night, and 4) white noise with Chebyschev filter centered at 4000Hz, which produced a waterfall-like sound. These sounds were designed according to ‘typical sounds perceived by tinnitus patients’ and represented auditory stimuli for a series of experiments within the framework of a larger project on top-down influences on sound perception in normal-hearing participants and patients suffering from tinnitus (German Research Foundation DFG, project number: 334628700). The order of the sounds, and thus the four different trials per task, was pseudo-randomized across participants. At the end of each tone/trial, participants were asked to rate on a continuous visual analogue scale how loud and how pleasant/unpleasant they perceived the sound. Continuous ratings were enabled by a sliding bar that could be manipulated left/right and up/down by button presses.

In the auditory and visual tasks, each trial contained eighteen modulations of the attended (i.e. auditory or visual) stimulus stream pseudo-randomly placed within the 100 seconds. For the auditory task, the modulations took the form of a 500ms reduction in stimulation volume that gradually reached its low of 50% of normal loudness after 250ms and then returned to normal volume within 250ms. In the visual task, the modulation was the fixation cross gradually changing its color from white to gray over 250ms, and then back to white within 250ms. In the visual task with auditory distraction, there were eight visual modulations to detect, which were identical to those in the visual condition. In addition to this, there were eighteen auditory modulations that consisted of superimposing a random choice of one of the other three background auditory stimuli on the already playing background sound for 500ms. Participants were instructed to ignore these audio modulations, and to rather focus on and respond to visual modulations. In the fourth task, participants were asked to count backwards from 500 while they heard the tones. After each tone, they were asked on what number they currently were to ensure that they followed the instructions. The experiment as well as the hearing test were programmed in Psychopy version 3 [43].

### Data acquisition

MEG data were acquired with a 4D Neuroimaging Magnes 3600 system with 248 data channels and 23 reference channels at a 678.2Hz sampling rate. Data were filtered online with a bandpass of 1-200 Hz, and corrected online for stationary, external magnetic noise with use of the reference channels. Structural MRIs were acquired using a high-resolution 3 T MRI-System (Siemens Magnetom Trio, Department of Neuroradiology, Universitätsklinikum Erlangen).

### Data analysis

#### Preprocessing

All analyses were performed in MNE Python v. 0.23 [44]. Data were first notch filtered at 50Hz and multiples thereof up to 200Hz, with an additional notch filter at 62Hz for a persistent noise source at this frequency, and then downsampled to 200Hz. Data were then manually inspected in order to identify bad channels and small sections with large bursts of noise. An ICA (Picard algorithm, [45]) was performed, limited to 60 components. Components which were clearly reflective of ocular muscle or cardiac noise were removed from the data. In addition, a reference channel ICA noise cleaning method was applied, which removes some sources of magnetic noise that are intermittent, and therefore not adequately compensated for by standard, online reference channel correction ([46]; “separate” algorithm). The cleaned data were then segmented into two-second-long epochs, which were the objects of all further analysis. Epochs that overlapped with a modulation or a motor response were excluded. Finally, some residual epochs with excessive noise, that were not picked up by the automatic methods, were excluded by hand upon visual inspection of the epoch power-spectra.

#### Source space and forward model

MRIs were segmented with Freesurfer [47–50]. For each individual MRI, a distributed cortical source space was created with ico5 spacing, resulting in 10242 sources per hemisphere. Single-shell, inner-skull (conductivity 0.3 S/m) boundary element models (BEM) were also constructed with Freesurfer, using the Watershed algorithm. These were then used to construct forward models.

#### Source localization and Parcellation

Before calculating connectivity, epochs were first source localized using sLORETA, with a regularization parameter of 1.

Source estimations were then consolidated into the Region Growing 70 parcellation described in [51]. This reduces the amount of data points to 35 in each hemisphere. Further, these 35 regions cover only areas of the cortex that are 1) robustly measurable at the scalp by MEG/EEG, and 2) have minimal localization-based cross-talk into or from other regions. In the case of connectivity estimation, consolidation from source points into regions was performed with the PCA flip method, which best preserves phase information across a range of different sources. The PCA flip method applies a Singular Value Decomposition to all sources within the region and selects the dominant component, which is then scaled to match the average power of the region, and if necessary flipped in polarity to match the polarity indicated by the dominant source orientation of the region. In the case of oscillatory power, a simple average was used for region consolidation.

#### Connectivity

Connectivity was calculated on the source-localized data in the theta (4-8Hz), low alpha (8-10Hz), high alpha (10-13Hz), beta (13-30Hz), and gamma (31-48Hz) bands using directed Phase Transfer Entropy (dPTE) [38]: Instantaneous phase of the source localized data was calculated with a morlet wavelet transform of 3, 5, 7, and 9 cycles for the respective frequency bands. For setting the delay and histogram bin width that are used in the dPTE calculations, we followed the practice of [39]: delay was set at (NxS)/(X_0_), where N is the number of samples, S is the number of signals (70 in this case, from 35 parcels per hemisphere), and X_0_ is the number of zero-crossings in all signals. Histogram bin width was set at e^0.626+0.4*ln(N-1)^, where N is the number of samples, and ln() is the natural logarithm.

#### Oscillatory power

Power in the high alpha (10-13Hz) band was localized using Dynamic Imaging of Coherent Sources (DICS) [52]. For each participant, DICS filters were produced from cross spectral densities calculated across all conditions with morlet wavelets of seven cycles, using only the real component for filter calculation. Power was then calculated separately for each individual epoch, and summarized into the regions of the Region Growing 70 parcellation by taking the mean of all values within a given region. DICS power was log transformed before statistics.

### Statistics

Statistical inference was performed with Linear Mixed Effects (LME) models, using the Python package Statsmodels 0.13.1. In all models we calculated, each individual epoch was a data point, and participant was a categorical random effect. Variables to compare different task conditions were then added as categorical fixed effects in a model comparison procedure.

#### Connectivity statistics with model comparisons

Our LME models testing connection strengths in the different frequency bands were designed to provide parameter estimates for changes in connectivity strength for different task conditions in relation to resting state. We always fit three separate models and compared their goodness of fit: a **null model,** which fit only an intercept, a **simple task model** which compared only between rest and task, without distinguishing different task conditions, and a **full task model**, where distinctions between task conditions were retained (audio task, visual task, visual task with distraction, backward counting). In the connectivity analysis for each frequency band, this fitting of the three models from null to full task model was performed for each of the 2415 possible connections between the 70 regions in a mass-univariate approach, using the Akaike Information Criterion (AIC) to assess goodness-of-fit and a Monte Carlo permutation to determine significance of differences.

The Akaike Information Criterion (AIC) is a measure of how well an LME model fits the data, weighted against how many parameters were used to fit the data, with lower AIC values indicating better fit given the number of parameters. As such, the AIC is a measure of goodness of fit that also favors model parsimony. This means that if for example the simple task model produces a lower AIC than the null model, this indicates that the explanatory power added by distinguishing between resting state and task was worth the reduced parsimony of adding a parameter to the model. The reduction of the AIC with improving models is referred to as the AIC delta, which is an interpretable quantity of model comparisons irrespective of the measurement units of variables in the models [53].

Like this, each connection was assessed as to whether the information provided by a distinction between rest and task was significantly better in explaining its strength over trials (simple task model) than assuming no difference (null model), and also, if distinguishing the task conditions yielded an additional significant explanatory improvement (full task model). Concretely, every connection where the null model was significantly worse at explaining the data than one or both of the task models was inferred to be a connection of potential interest. Connections where no model performed significantly better than null were inferred to be unaffected by the experiment, and not analyzed further. Then, out of the connections of potential interest, those connections where the simple task model performed significantly better than the null model, but the full task model did not perform significantly better than the simple task model were considered best explained by the simple task model. Finally, connections where the full task model performed significantly better than the simple task model were considered best explained by the full task model.

To decide which reduction of AIC value from null model to simple task model, or from simple task model to full task model was deemed ‘significant’ in increasing explanatory power, and to control for the multiple comparisons in our mass-univariate approach at the same time, we used a permutation approach that solved both problems. We permuted the condition labels across subjects’ epochs 1024 times and fit the simple and full task models to the permuted data for all connections, collecting their respective AICs. By subtracting the AIC of the null model of each connection from the respective AICs of the simple task model of the permuted data, a surrogate distribution of AIC reductions (AIC deltas) expected under the null hypothesis was derived. For each connection, the greatest AIC reduction across the 1024 permutations was noted, and the significance threshold for the AIC deltas for simple task model vs. null model was set at the 0.05/2 quantile of the maximal reductions across all 2415 connections. In the same vein, the significance threshold for the AIC deltas showing improvement for the full task vs. simple task model was derived by subtracting the average permuted simple task model AICs from the AICs of the full task model of the permuted data, collecting the maximal AIC delta (reduction) for each connection across permutations, and calculating the 0.05/2 quantile of the maximal reductions across all connections. This yielded an AIC delta significance threshold of 14.97 for assessing simple task vs. null model, and of 19.88 for full task vs. simple task model comparisons. Given that AIC deltas of 2 are seen as meaningful and of >6 as showing strong explanatory improvement, our principled approach and calculation clearly derived strict criteria for assessing meaningful connectivity changes.

#### Post-hoc test on motor/parietal – sensory connections

Because the three tasks involving motor responses differed in which modality required attention to detect modulations– namely audio, visual, or visual with auditory distractions, and the significant changes in connectivity between the motor/parietal hub and sensory cortices in the high alpha band encompassed both auditory and visual cortices (see Results), we performed a post-hoc statistical test with a linear mixed effects (LME) model restricted to the connection between the motor/parietal hub and the primary auditory cortex (A1), and the connection between the motor/parietal hub and the primary visual cortex (V1). Like with our other models, data points were individual epochs, and participant was entered as categorical random effect. Our dependent variable was the sum of zero-centered dPTE values (subtracting 0.5) between the motor/parietal hub (comprising 4 parcels) and the relevant primary cortex in the left hemisphere. Categorical fixed effects included condition/task (resting state, audio task, visual task, and visual task with distraction) and primary cortex destination (A1 and V1). Parameters for the interaction between the condition and destination effects were also estimated.

#### Connectivity-power influences in the alpha band

We also directly explored the relationship between dPTE and oscillatory power in the high alpha band, focusing on its changes from rest to task(s) for connections from the motor/parietal hub to the primary sensory cortices (A1, V1) and their influence on alpha power in those regions (A1, V1). After calculating high alpha power with DICS for each region (see Oscillatory power above), this relationship was tested with two linear mixed effects models, one for each primary sensory cortex (A1, V1). Here, on a trial-by-trial basis, local log-transformed alpha power was analyzed as the dependent variable, with the dPTE calculated from the combined motor-parietal hub to the respective area (A1 or V1) modeled as independent, fixed effects variable, together with the categorical factor task (rest, visual, audio, visual with distraction) and their interactions, and participant modeled as random effect.

#### Potential SNR/dPTE confounds

Past work has raised the possibility that SNR differences in oscillatory power can affect instantaneous phase estimation, which could in turn cause spurious connectivity findings with any method that makes use of phase [54], which would include dPTE. Under this account, the observed dPTE changes we see could be simple by-products of alpha power changes. In order to test for this confound, we carried out single-trial, linear mixed model comparisons for the left parietal hub in the high alpha band - the region and frequency band where we saw the most prominent, theoretically important changes to connectivity by condition. Specifically, we compared the explantory power of models which 1) used only condition to explain dPTE values in the left parietal hub, 2) used only log power to explain dPTE, and 3) used both condition and log power. If dPTE is in fact only an artifact of varying SNR (power), we would not expect the condition + power model to have more explanatory power than the power only model, as assessed by the models’ AIC and AICdeltas.

## Results

The first part of our study is aimed at describing the general patterns of directed connectivity via directed phase-transfer entropy (dPTE) of oscillations in different frequency bands between 70 regions of a parcellation optimized to study connectivity in the MEG, in comparison of an eyes-open resting state and the task-states induced by our set of attention tasks. We report here separately for the five analyzed frequency bands (theta, low alpha, high alpha, beta, and gamma) those connections that change their strength or direction reliably from resting state, as identified by the linear mixed effects and permutation approach described in the methods section. A majority of significant changes was best explained by a simple task model describing a shift to a ‘general task state’ across all our sub-tasks. In turn, we focus on the high alpha band (10-13 Hz), where we see the strongest qualitative changes from resting state, to investigate how connectivity changes from a motor/parietal hub to the primary sensory cortices are associated with the different attention tasks as well as their relationship with the modulation of sensory alpha power.

### Connectivity patterns and changes with task

The resting state connectivity patterns and significant changes that relate to a general task state (‘general task’) or specific attention tasks (‘audio’, ‘visual’, ‘visual with distraction’, ‘counting backwards’) are summarized and displayed in Figures 1 to 3 for the lower frequency bands (see Fig. S1 for the beta and gamma bands). In each Figure, section A) displays the top 150 connections in the respective frequency band during resting state, indicating the direction of flow from source (red) to target (blue). The B) sections, then, visualize the full resting state connectivity pattern in matrix form (top left), together with the significant changes during tasks, as revealed by the linear mixed effect ‘simple task models’ and ‘full task models’. Each resting state matrix plot in section B) shows the directionality and strength of all connections between the 70 regions at rest, grouping the fine lines of single regions into broader color-coded areas on the axes (occipital, parietal, central, frontal) divided by hemisphere (lh, rh; coded by lighter or darker shade) to improve readability of general patterns. Significant connection changes that can be attributed to a general task state as estimated by the simple task model are displayed in the matrix ‘general task’ (bottom right). Any additional changes that significantly improve the model when differentiating between the four attention tasks, as estimated by the full task model, are shown in the single task matrices for ‘audio task’, ‘visual task’, ‘visual with distraction’, and ‘counting backwards’. For reading and interpreting the matrices correctly, it is important to keep the following information in mind about scaling and color coding. As dPTE values express both the dominant directionality of a connection, as well as its strength, we centered them around 0 (subtracting 0.5) and used a bidirectional color scale, whereby darker shades of red denote a stronger influence or flow from region X to region Y, and darker shades of blue a stronger influence or flow from region Y to region X. As the matrices for the single task and general task changes display the significant parameters of the respective linear mixed effects model, the values, i.e. colors, here have to be interpreted with respect to the resting state pattern, as they indicate changes from rest, not absolute dPTE during the respective task. This means that, e.g. a light blue dPTE value in a task matrix for a connection that had a medium red value in the resting state matrix displays a reduction of the X-to-Y flow seen at rest, whereas a red value for the same connection would mean a further increase in X-to-Y connectivity, while a deep blue could indicate a flow reversal with Y-to-X influence becoming slightly more dominant during task.

**Figure 1:**
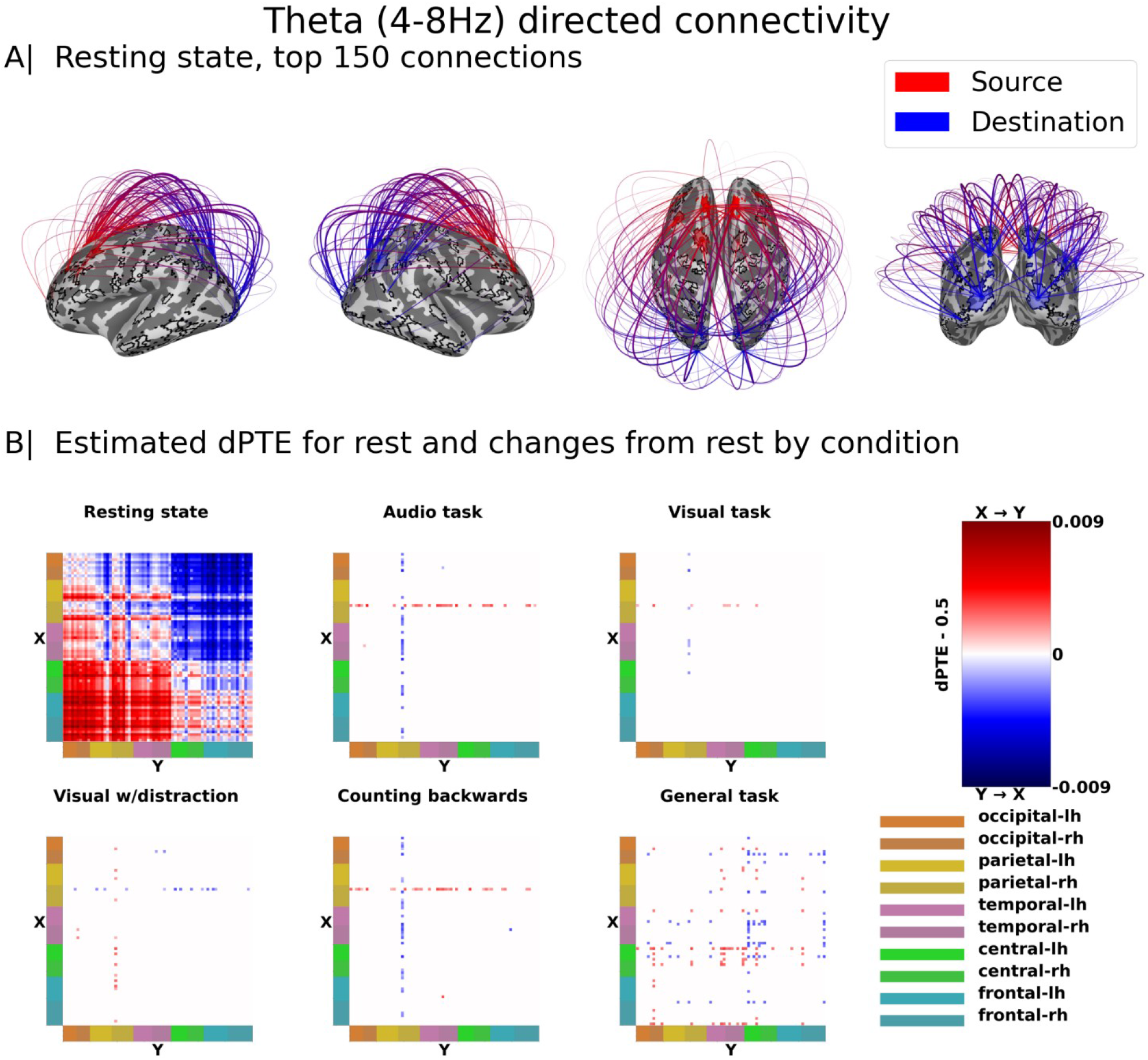
directed Phase Transfer Entropy (dPTE) in the theta band (4-8 Hz). **A)** The top 150 strongest connections during resting state. Information movement from source to destination is indicated by red to blue. B) Matrices displaying dPTE connection strengths at rest, and significant divergences from rest by condition estimated with linear mixed effects models. Values in the resting state matrix are zero-centered dPTE values (subtracting 0.5) and colors thus show both direction and strength (with darker shades of red indicating stronger X to Y flow, and darker shades of blue stronger Y to X flow), corresponding to the model intercepts. The condition matrices of the different tasks display the further model parameters, i.e. significant estimated changes from that baseline. Thus, values and colors in the condition matrices have to be interpreted with respect to the same connection at rest (e.g. if a connection has strong X to Y flow at rest (red), negative values (blue) of lesser or similar amount show a reduction of that outflow).

Figure 4 A-E provides an additional visualization and summary of the main patterns of change of the dPTE in the theta and low and high alpha bands during the tasks. It displays the significantly changing connections on an inflated brain with the parcellation used, that correspond to the respective dots and lines from the matrix plots in Figures 1-3. Colors in the brain connectivity plots, thus, have to be interpreted with respect to the resting state flow, just like the change the LME change parameters in the matrices of Figures 1-3. Additional bar charts visualize the changes in summed dPTE values of the most prominent/affected regions or hubs for each frequency band. Substantial changes with pronounced differences between sub-tasks were primarily seen in the high alpha band (10-13 Hz), Fig. 4 D-E, whereas other change patterns seem more indicative of a shift into a general task state, Fig. 4 A-C.

### Theta band (4-8 Hz)

In resting state, there is a general anterior to posterior transfer of information in the theta band, replicating [39] (Fig. 1A). During tasks, there is an increase in the already existing outflow from the left motor cortex primarily to the temporal lobes, and less so to a broader range of areas. The right frontal pole also increases its outflow to a lesser extent (Figs. 1B, 4A). In the right dorsal parietal cortex, there is a reduction of information inflow particularly in the backwards counting (internal attention) and auditory conditions, while inflow increases in the visual task with auditory distractions.

### Low alpha (8-10 Hz)

Information in the low alpha band (8-10 Hz) flows from posterior to anterior areas at rest, replicating [39] (Fig. 2A). During all tasks, the major pattern of change in connectivity shows an increase of the existing information flow from several occipital and temporal areas, including the sensory cortices, to the parietal, central and frontal lobes (Figs. 2B, 4B) One right dorsal parietal region also increases its flow to several central and frontal regions, parallel to some minor flow increases from frontal to central sites. Overall, the statistical ‘simple task model’ was preferred for the majority of connectivity changes, as seen in the dominant pattern in the ‘general task’ matrix (Fig. 2B), suggesting that the low alpha band dPTE here primarily describes a shift into a general task state. Some additional differences between single tasks, i.e. significant changes in connection strength favoring the ‘full task model’, were also observed, however. The left mid-temporal lobe increased its outflow to parietal and frontal regions during all three tasks requiring motor responses (i.e. except backwards counting); in the visual and visual with distraction tasks it also increased its flow to occipital and right temporal regions. The right V1 increased its existing outflow to several frontal regions from resting state in all tasks except the visual with distraction task, with audio and backwards counting conditions showing stronger increases than the visual task, and additionally increased flow to left temporal regions. A further, left occipital region increased its flow to several parietal and a few frontal regions during both visual tasks (without/with distraction), while receiving more local inflow from other occipital sites in the audio and the backwards counting task.

**Figure 2:**
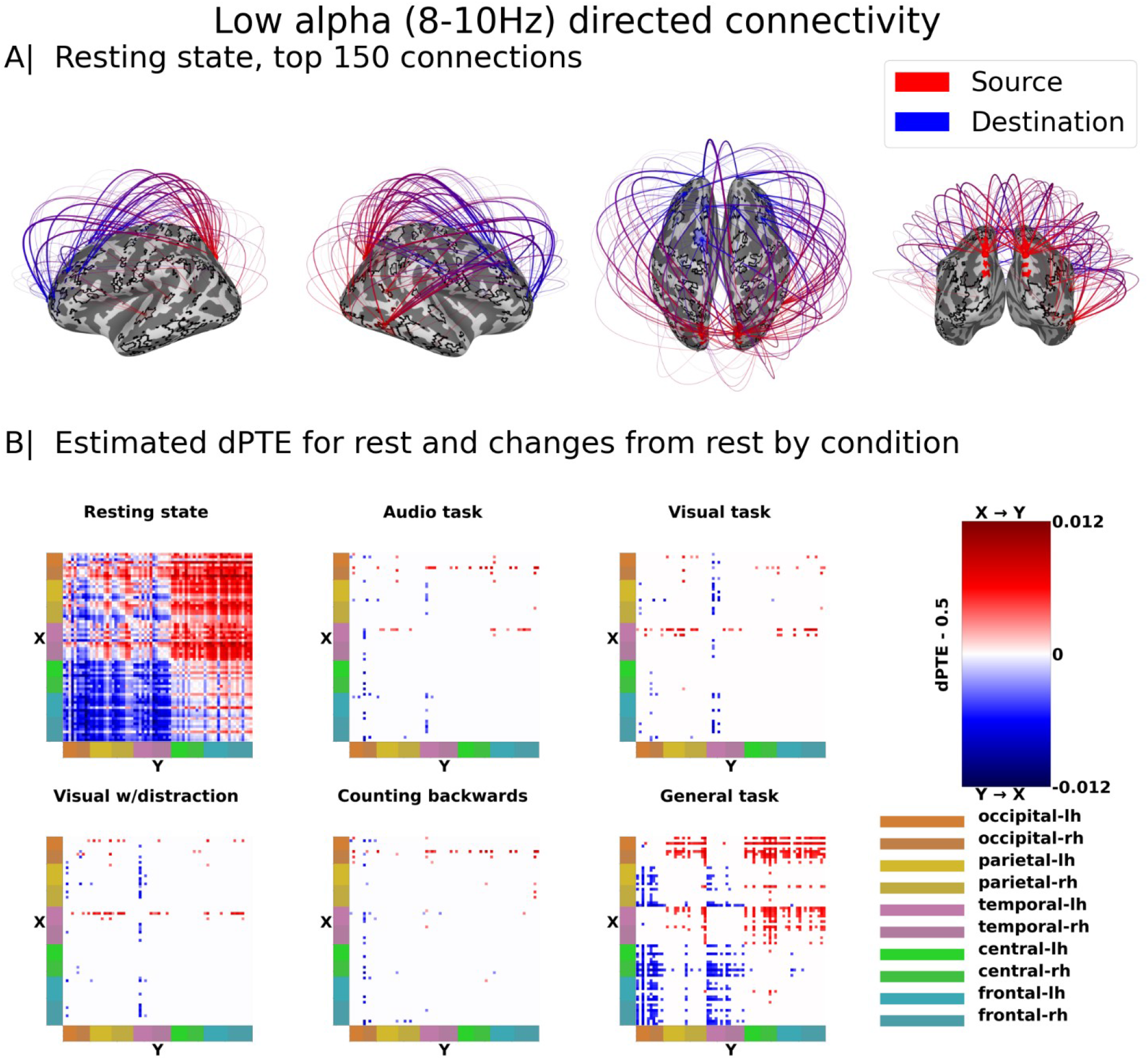
directed Phase Transfer Entropy in the low alpha band (8-10 Hz). A) The top 150 strongest connections during resting state. Information movement from source to destination is indicated by red to blue. B) Matrices displaying dPTE connection strengths at rest, and significant divergences from rest by condition estimated with linear mixed effects models. Values in the resting state matrix are zero-centered dPTE values (subtracting 0.5) and colors thus show both direction and strength (with darker shades of red indicating stronger X to Y flow, and darker shades of blue stronger Y to X flow), corresponding to the model intercepts. The condition matrices of the different tasks display the further model parameters, i.e. significant estimated changes from that baseline. Thus, values and colors in the condition matrices have to be interpreted with respect to the same connection at rest (e.g. if a connection has strong X to Y flow at rest (red), negative values (blue) of lesser or similar amount show a reduction of that outflow).

### High alpha (10-13 Hz)

In the high alpha band (10-13 Hz), the general resting state information flow was also primarily from occipital and parietal areas to frontal areas, but with temporal areas also receiving generally strong inflows (Fig. 3A), again replicating [39]. Another resting state pattern specific to this frequency band is a strong information flow from the motor and parietal cortices to occipital and temporal regions, including the primary sensory (visual and auditory) cortices. In the general task state, there was an overall reduction in the outflow from a left parietal hub (left posterior parietal cortex) to the entire cortex. This reduction was most pronounced in connections to the temporal and occipital lobes, and was somewhat weaker in connections to the frontal lobes (Figs. 3B, 4C). During the three attention tasks which required a motor response (audio, visual, visual with distraction), a left hemispheric motor/parietal hub, encompassing again left posterior parietal cortex and, additionally, left primary motor and somatosensory cortex, decreased its outflow to most areas of the cortex, mostly nullifying the resting state outflow (Fig. 3B, 4D). During the task involving backwards counting (internal attention) there were modest increases in already existing outflow from bilateral temporo-parietal junctions to temporal, central, and frontal areas, albeit with left hemispheric dominance (Fig. 3B, 4E).

**Figure 3:**
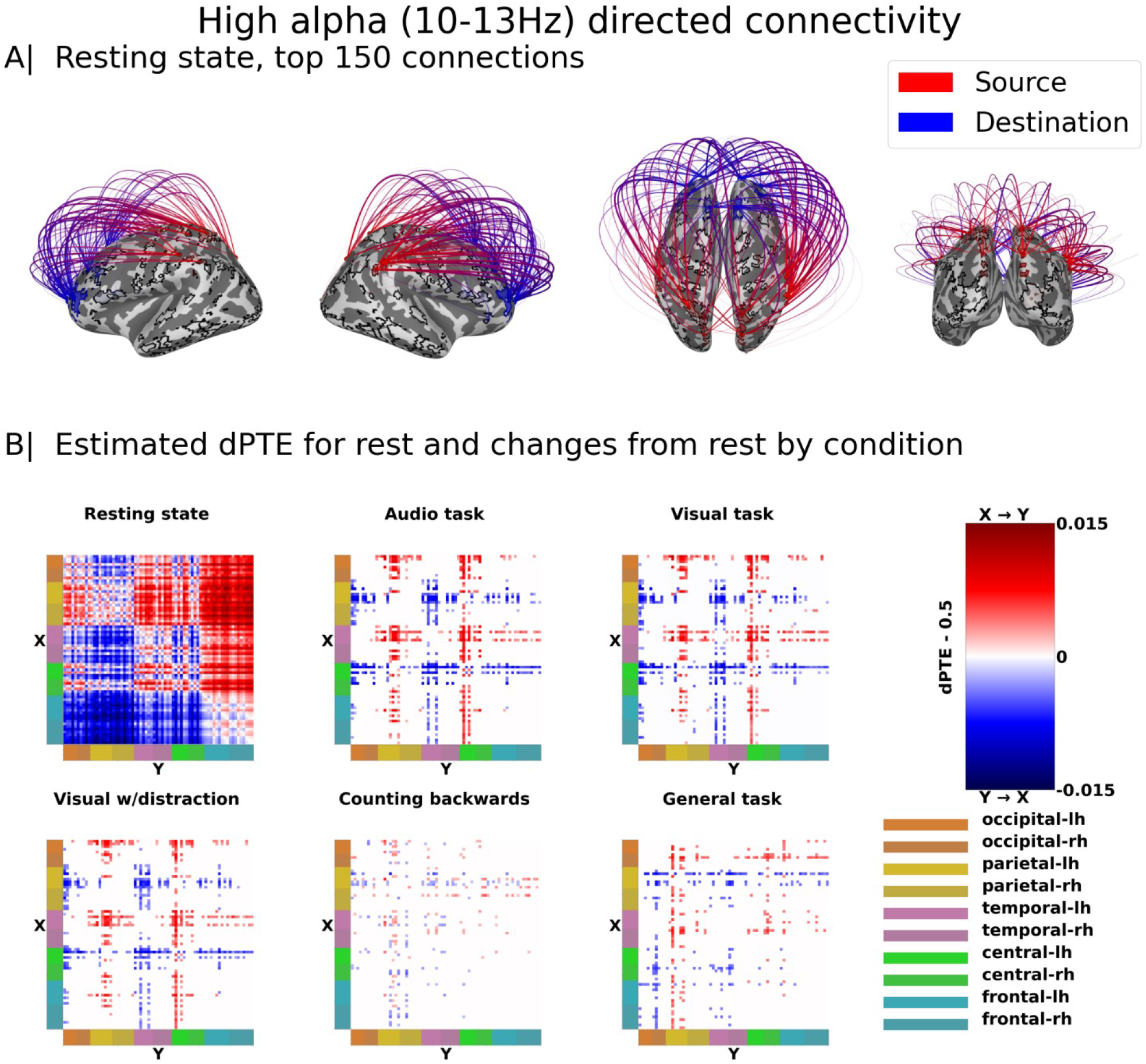
directed Phase Transfer Entropy in the high alpha band (10-13Hz). A) The top 150 strongest connections during resting state. Information movement from source to destination is indicated by red to blue. B) Matrices displaying dPTE connection strengths at rest, and significant divergences from rest by condition estimated with linear mixed effects models. Values in the resting state matrix are zero-centered dPTE values (subtracting 0.5) and colors thus show both direction and strength (with darker shades of red indicating stronger X to Y flow, and darker shades of blue stronger Y to X flow), corresponding to the model intercepts. The condition matrices of the different tasks display the further model parameters, i.e. significant estimated changes from that baseline. Thus, values and colors in the condition matrices have to be interpreted with respect to the same connection at rest (e.g. if a connection has strong X to Y flow at rest (red), negative values (blue) of lesser or similar amount show a reduction of that outflow).

**Figure 4:**
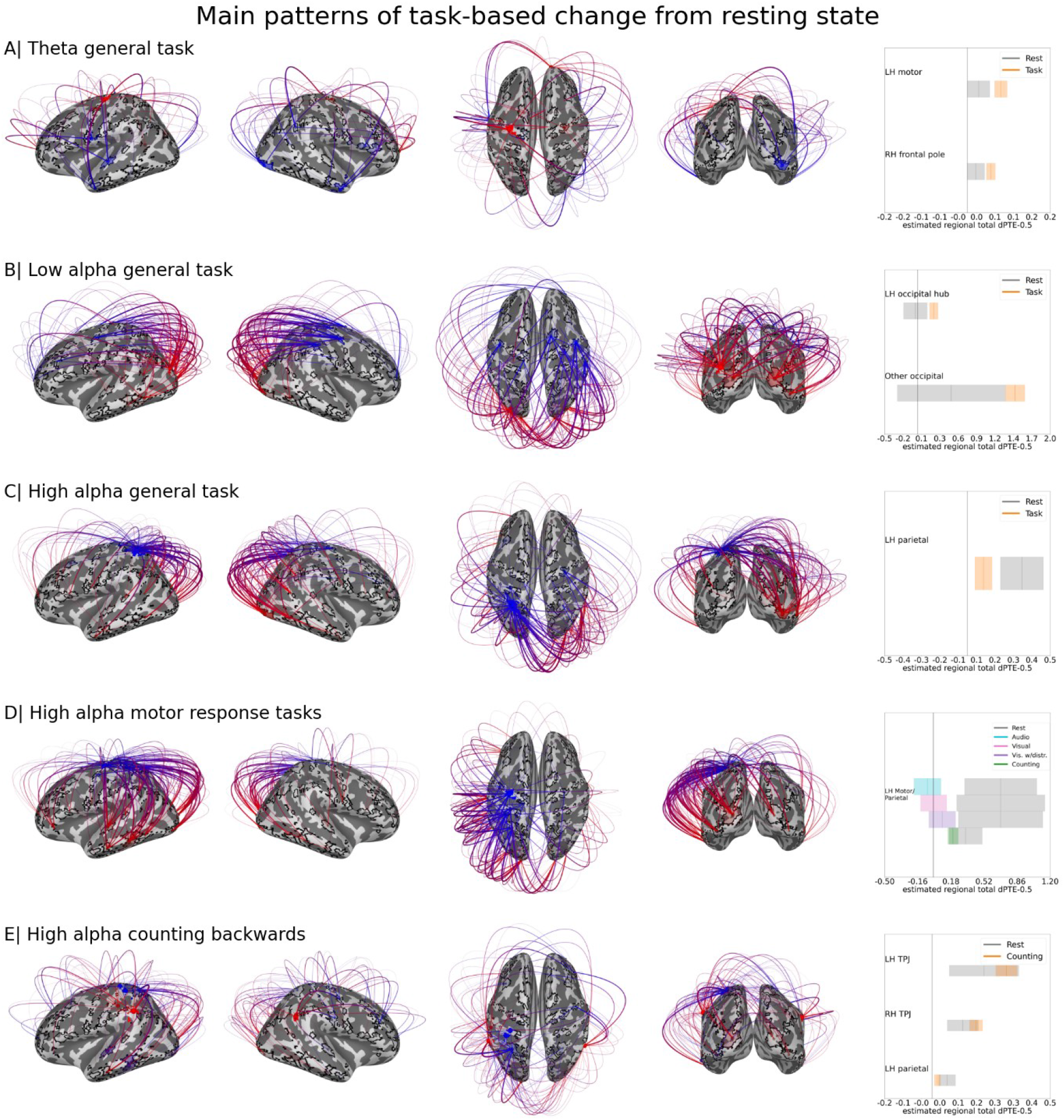
Main patterns of dPTE changes from resting state to task state. A) Changes to theta connectivity moving from resting state to any task for hubs in the left motor region and the right frontal pole. B) Changes to low alpha connectivity moving from resting state to any task for a left occipital hub, and the rest of the occipital lobe excluding the former hub. C) Changes to high alpha connectivity moving from resting state to any task in a left parietal hub. D) Changes to high alpha connectivity moving from resting state to all four different tasks in a left motor/parietal hub. E) Changes to high alpha connectivity moving from resting state to the backwards counting task in the left and right temporoparietal junctions and a left parietal hub. Brain connectivity plots on the left show estimated connectivity changes from resting state, restricted to the most prominent hubs. Bar plots on the right show the resting state total dPTE (grey bars) and change by task(s) (colored bars) with 95% confidence intervals, as estimated by a LME. Total dPTE here means the zero-centered sum of a hub’s connectivity to all other regions. The bar plots also help interpret the colors of flow changes in the brain connectivity plots (blue/red), i.e. outflow increases of red hubs in A) and B) vs. outflow decreases of blue hubs in C) and D).

### Beta (13-30 Hz) and Gamma (31-48 Hz)

In the higher frequency bands, beta (13-30 Hz) and gamma (31-48 Hz), the overall divergences of the dPTE values from neutral were about an order of magnitude lower than those seen in the theta and alpha bands. In the beta band during resting state, there was a general flow of information primarily from bilateral parietal areas to temporal, central, and frontal areas. In the gamma band during resting state the bilateral temporo-parietal areas received information mostly from single hubs in the bilateral occipital, temporal, frontal, and fronto-central lobes. In each of the two frequency bands, only one connection out of 2415 was significantly different between rest and task; we leave them uninterpreted here. Resting state patterns are depicted in Figure S1.

### Task-specific connectivity changes and relation to alpha power

While a substantial amount of the observed connectivity changes from resting state to our set of attention tasks is most likely related to a switch into a general task state, a specific set of connections in the high alpha band (10-13 Hz) stands out, that seems to be pertinent to qualitative changes related to the specific tasks applied here. Most pronounced in this respect are the flow changes between resting state and the external attention tasks, i.e. those including a motor response between a left-hemispheric motor/parietal hub and left occipital and temporal areas. As we had expected changes in alpha band connectivity to the task-relevant primary sensory areas, we carried out post-hoc tests on connection strengths across these three tasks (audio, visual, visual with auditory distraction) between the left motor/parietal hub (M/P) and the left primary auditory cortex (A1) and the left motor/parietal hub (M/P) and the left primary visual cortex (V1) (see Methods). The parameter estimates of the respective linear mixed effects model (LME) along with their 95% confidence intervals are depicted in Figure 5 (left side) - all estimated parameters were statistically significant at *p* < .005. In the motor/parietal(M/P)-to-A1 connection, the strong resting state information flow out from M/P is significantly reduced for all experimental tasks. This reduction is numerically strongest for the auditory task, but, given the degree of overlap of confidence intervals, is merely suggestive of a functional interpretation. In the motor/parietal(M/P)-to-V1 connection, the prominent resting state information flow from M/P is also significantly reduced for all experimental tasks. Again, the reduction here is numerically strongest for the visual task, suggesting a functional relevance. The same cautionary interpretation given overlapping confidence intervals between tasks as for the auditory cortex applies here. In summary, each primary sensory cortex shows the greatest inflow reduction from the motor/parietal hub when the associated modality was task-relevant (attended) and without distraction. Moreover, we see a slight mean reversal of information flow from the auditory cortex to the motor/parietal hub when task-relevant. This pattern is a bit less clear/interpretable for the visual cortex, where both visual and audio task seem to show a slight flow reversal (albeit stronger for visual). In both regions, the inflow reduction from M/P is weakest in the visual task with distraction.

**Figure 5:**
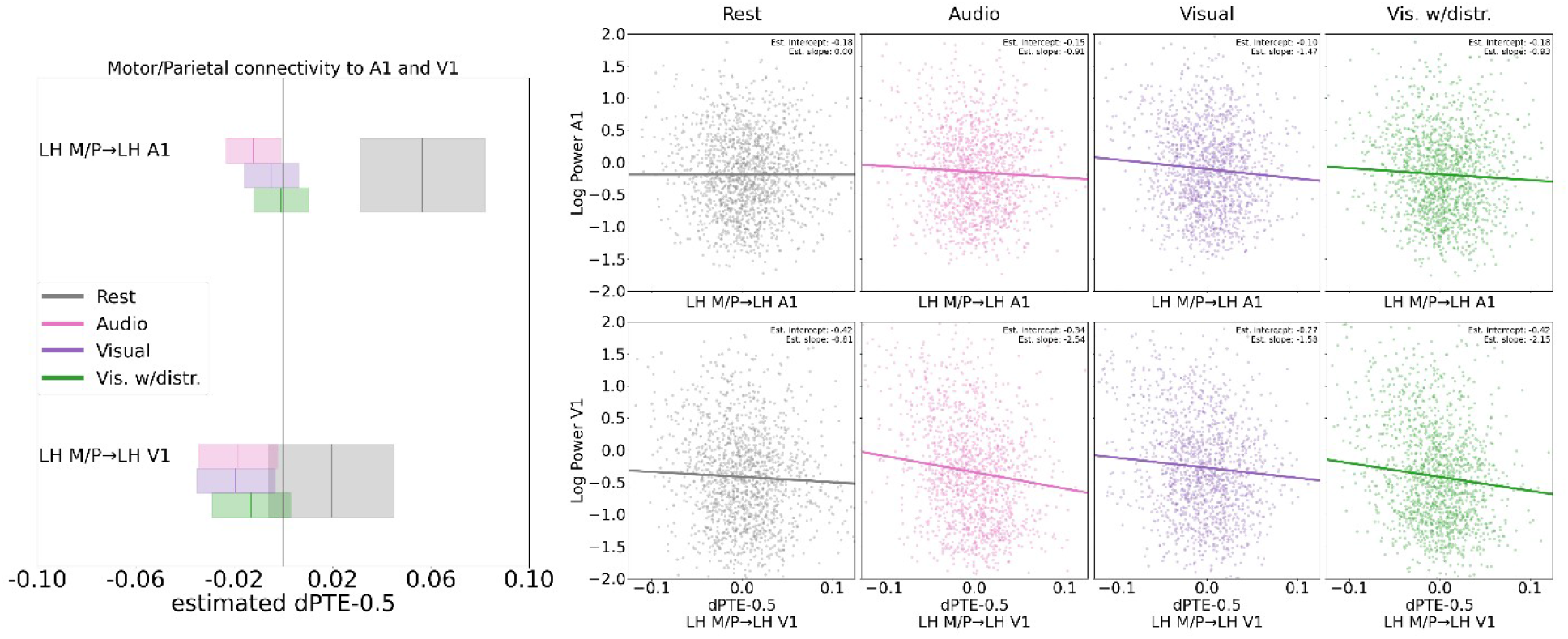
**Boxplots, left:** Connectivity changes between the left motor/parietal hub and left primary auditory cortex (top) and left primary visual cortex (bottom) during attention tasks with motor response from a resting state baseline. Boxes represent 95% confidence intervals and vertical lines within boxes represent estimated dPTE values, in both cases calculated from a linear mixed effects model. Connectivity from the left motor-parietal hub to the auditory cortex is reduced for all tasks involving motor responses, strongest however for the auditory task (pink rectangle). In contrast, connectivity from the left motor-parietal hub to the visual cortex is reduced most for the visual task (purple rectangle). **Scatterplots, right:** Trial by trial relationship of log-transformed, high alpha band oscillatory power in primary auditory (top)/visual (bottom) sensory cortices and dPTE from the motor/parietal hub to the auditory/visual cortex; positive dPTE values indicate flow from M/P hub to sensory cortex and negative indicate flow in the reverse direction. Top row shows M/P hub to A1, and bottom row shows M/P hub to V1. Columns show the relationship across resting state and the three motor response tasks. The intercept and slope were calculated by combining significant parameters from linear mixed models that had either A1 or V1 log power as a dependent variable, and dPTE from M/P to A1 or V1 and Condition as independent variables. In A1, the slope is steepest for the visual task, whereas in V1, the slope is steepest for the audio task.

Finally, we wanted to test whether these task-specific changes in alpha band connectivity are indeed associated with local alpha power levels in the sensory cortices, supporting our hypothesis that connectivity or changes in information flow might drive attention-related power modulations. Two further LME models were calculated to this end, modeling trial-by-trial log power in each primary sensory cortex (A1, V1) by the dPTE flow from the motor/parietal hub (M/P) to this region and by experimental condition (rest, audio, visual, visual with auditory distraction). Figure 5 (right side) displays the results for A1 (top row) and V1 (bottom row), showing scatterplots of the dPTE-power relationship together with their estimated intercepts and slopes, i.e. log power and change by dPTE per condition.

General task-related increases in local alpha power are seen in both primary sensory areas for both the audio and visual task, but not the visual task with distraction. Log alpha power in A1 increases by .036 for the audio task (*p* < .01) and by .079 for the visual task (*p* < .001), while log alpha power in V1 increases by .071 for the audio task (*p* < .01) and by .141 for the visual task (*p* < .001), and mean power levels in the visual task with distraction stay the same as in resting state (*p* = .79 and *p* = .12 for A1 and V1 respectively). Thus, in the simple tasks without distraction, sensory alpha power increases significantly for both the attended and task-irrelevant modality – and generally more so for the visual task and in visual cortex.

In contrast, the trial-based alpha power shows a much clearer picture of task-related modulations when seen in relationship to the dPTE flow from/to the motor/parietal hub (M/P). In the auditory cortex, no significant dPTE to power association is visible at rest (*p* = .15), but during the three attention tasks with motor responses, the flow from M/P to A1 has a significant negative relationship with log alpha power in A1 – with *Coef* = −0.91, *p* <.01 during the audio task, *Coef* = − 0.93, *p* <.01 during the visual task with auditory distraction, and *Coef* = −1.47, *p* <.001 during the visual task. Knowing that during these tasks mean M/P-to-A1 alpha dPTE reduces (Fig. 5 left side), these negative relations thus indicate trial-wise local A1 alpha power increases with these reductions – strongest for the visual task. Also, even though there is no main effect on power showing a general power increase for the visual task with auditory distraction, the model suggests that there are indeed alpha power increases in line with dPTE flow reductions on a trial-by-trial basis. In the primary visual cortex, there is already a significant negative relation of M/P dPTE outflow and V1 alpha power at rest, *Coef* = −0.81, *p* <.005, which is then intensified during all three attention tasks – by *Coef* = −0.77, *p* <.05 during the visual task, *Coef* = −1.34, *p* <.001 during the visual task with auditory distraction, and by *Coef* = −1.74, *p* <.001 during the audio task. As for A1, the negative relationships can be interpreted as trial-wise alpha power increases related to the known M/P outflow reductions during these tasks. Again, the trial-wise model also shows a clear dPTE-power relationship for the visual with distraction task, where no global power increases were observed in form of a main effect.

In summary, in both primary sensory cortices, alpha power has a significant negative relationship with inflow from M/P during all tasks, i.e. local power increases with the reported M/P outflow reductions (as seen in Fig. 5, left side), that is significantly different from resting state. Moreover, the slope of this relationship is flattest for the attended or task-relevant modality (audio in A1, visual in V1), intermediate for the visual task with auditory distractions in both A1 and V1, and steepest for the task-irrelevant modality (visual in A1, audio in V1), showing that trial-by-trial M/P alpha dPTE outflow reductions are associated with alpha power increases in a task-sensitive way.

To rule out a possible SNR/dPTE confound, we used linear mixed model AIC comparisons on single trial high alpha dPTE and log power values in the left parietal hub in order to ascertain to what degree condition-based changes in dPTE actually occur independently of power changes - power here being a proxy for SNR, which has been suggested may cause artifactual connectivity (see Methods and Discussion). The model which explained dPTE with power alone had more explanatory power than the model which used condition alone (AICs: -24681 and -24616, respectively). The model which used both power and condition to explain dPTE however performed better than the power alone model (AIC -24703, with an AIC delta of 22 to the power-only model), indicating that changes to condition affect dPTE values independently of power (SNR) differences.

## Discussion

Applying directed phase transfer entropy on source-localized magnetoencephalography we were able to show that the recruitment and integration of processing resources during the performance of four different attention tasks is mediated by a task- and region-specific modulation of alpha band connectivity. In a first step, we replicated the basic oscillatory network patterns at rest found in past studies [39] where in resting state information flows from anterior to posterior regions in the theta band, and from posterior to anterior regions in the alpha and beta bands. By introducing a set of related attention tasks, we could reveal a significant, qualitative change to the resting state pattern specific to the upper alpha band (10-13Hz), whereas in the theta and lower alpha band the information flow during attentional tasks tended to increase in the same direction as the resting state information flow. This qualitative change in the upper alpha band manifested as a decrease of the strong information outflow from the left posterior parietal and the left motor cortex to a majority of brain regions in the switch from resting state to task. While motor/parietal outflow was reduced toward many regions throughout the cortex, the most focused reductions were seen in connection to the task-relevant sensory cortices, namely temporal (auditory) and occipital (visual) areas, and somewhat less so to frontal areas. Moreover, these connections exhibited a high degree of qualitative differences between the single attention sub tasks, suggesting that they are functional for task-related differences. Crucially, the left motor cortex resting state information outflow was practically nullified when a motor response was required, but not during backwards counting, when no motor reaction was necessary. Backwards counting mostly increased existing connectivity in the bilateral temporo-parietal junction.

Between the three tasks requiring motor responses, there was a (statistically marginal) double disassociation in the flow reduction from the left motor/parietal (M/P) hub to the sensory cortices by attention modality, such that the flow reduction from left M/P to auditory cortex was strongest under auditory attention, and the flow reduction from left M/P to primary visual cortex was strongest under visual attention.

Finally, we found a reliable negative association between alpha-band dPTE outflow from M/P to the task-relevant sensory cortices and sensory alpha power, meaning alpha power in auditory and visual cortex was rising in relation to the flow reductions from M/P. There was a further double dissociation by attention modality, that appeared in this association more clearly than in the dPTE alone. Namely, the association between the alpha flow reduction from M/P to the sensory cortices and the increase of alpha power within these was strongest in auditory cortex for the visual task and strongest in visual cortex for the auditory task, i.e. each sensory region showed the strongest association when its suppression was task-relevant. When its modality (and not suppression) was task-relevant, however, each sensory cortex showed the weakest association with alpha power increases; while an intermediate-strength association was observed for the visual task with auditory distractions, where both modalities competed.

We discuss in the following, how these findings could be integrated into a theory of task-optimized neural resource management in the alpha band.

### Alpha flow modulations occur in a region and task specific manner setting up an ‘optimal task state’

Alpha band connectivity was systematically modulated from resting state in a region- and task-specific manner with a set of related attention tasks. The left dorsal parietal cortex, which strongly reduced its alpha outflow to the visual and auditory cortices processing the incoming visual and auditory stimuli participants were attending to, has been associated with attentional processes and top-down mediated gating of information processing [21, 55–57]. Our results are consistent with a role of the dorsal parietal cortex as an important hub for the distribution of attentional resources toward lower-level stimuli-processing areas.

In addition to the left dorsal parietal cortex, left motor cortex also markedly reduced its strong alpha outflow towards the visual and auditory cortices. Crucially, this was only the case when right-handed motor responses were required and not during the backwards counting task. This suggests that the motor cortex is specifically facilitating processing paths between itself and the relevant visual and auditory cortices associated with the required motor response.

That sensory areas might be modulated by the motor cortex based on the task-specific intended action, has previously been argued by Gallivan and colleagues [58, 59], who showed in their studies that the movement effector (specific limb) could be reliably decoded during motor preparation from both the visual and auditory primary sensory cortices. In a related fashion, we propose that in our experiment the motor cortex relevant for the intended action (right-hand button press) facilitates information transfer from the relevant sensory cortices to allow a rapid and appropriate output response. Gale et al. [59] also report that the decoding of motor effector can be achieved from bilateral auditory cortex during motor execution, but only from contralateral auditory cortex during motor planning, and suggest that motor planning and preparation employ an “effector-related global gating mechanism”. Our alpha connectivity modulations, which are also observed during time segments of motor preparation or planning (waiting for the appropriate sensory signals for a motor response) and mainly in the contralateral hemisphere to the executing limb (right hand), seem to represent such a gating setup for the task. Remembering that in the current study we are exclusively observing time segments of an ongoing audio-visual stimulation in which participants are preparing themselves to detect and respond to tiny modulations within the visual or auditory stream (since we cut out the time segments containing the actual modulations of the stimulus material and the button presses), the observed general and task-specific reductions of motor/parietal-to-sensory alpha flow could be interpreted as preparation processes supporting and maintaining an optimal state by opening and adjusting the gates of pathways specific and relevant to the current task. That the functional attention-related alpha connectivity changes in the outflow from the dorsal parietal cortex to the task-relevant auditory and visual cortices are also most pronounced in the left hemisphere in our data also fits into this interpretation, as they would be connected to and integrated with the left motor cortex for this task-optimized flow setup.

Summarizing again, parietal-to-sensory alpha flow is reduced in all tasks (having the same audio-visual stimulation), but much more so in the tasks requiring external attention including detection of modulations and a motor response than during internal attention/backwards counting. Motor-to-sensory alpha flow is strongly reduced only for the tasks requiring motor responses and not for backwards counting. These changes seem reflective of the setup of general task states for optimal processing, especially in the case of the three motor-response tasks, connecting the appropriate input (auditory, visual) and integration/output areas (parietal and motor). Under this account, both auditory and visual processing pathways are ‘opened’ by the outflow reduction of high alpha to the two sensory cortices in all motor-response tasks, allowing the most efficient information flow for the task at hand, channeling attention to relevant changes in the stimuli and enabling a quick motor response. This account could be investigated in further experiments by comparing similar attention tasks with and without required motor responses and with varying response hands in a more systematic way.

### Internal processing and the temporo-parietal junction

During silent backwards counting, we found bilateral increases in the alpha-band dPTE from the temporo-parietal junction (TPJ) primarily to frontal areas and secondarily to temporal and occipital areas. Backwards counting was mainly included as a control task directing attention away from both the auditory and visual sensory input, rather than to systematically explore the neural correlates of counting. Nevertheless, as the TPJ has both been associated with enumeration and number processing in the brain [60–62] and has been consistently implicated in introspection and internal processes ([63, 64] for recent review), its prominent role in the connectivity changes observed here with silent counting seem to match these ascribed functions.

### Reduced alpha flow opens task-relevant processing paths

A down-regulation of alpha oscillations associated with disinhibition and increased information processing is reminiscent of the gating-by-inhibition literature [2]. Modulations of ongoing alpha oscillations in sensory brain regions modulate local processing of e.g. visual and/or auditory stimuli during attentional tasks [9, 16, 65–69]. We focused here on *alpha connectivity in the whole brain* to understand better how alpha oscillations contribute to the distribution of processing resources in the brain, as opposed to alpha power levels only in specific brain regions. The task- and region-specific alpha flow reduction we found here by a left motor/parietal hub could reflect a mechanism of ‘opening processing gates’, which prioritizes task-relevant pathways and facilitates information transfer and integration from the relevant sensory cortices. Moreover, the general motor/parietal (M/P) flow reduction to the auditory and visual sensory cortices was further modulated in line with the specific target-modality of the external attention tasks, showing the strongest flow reductions to each sensory cortex when it supported the target modality (auditory or visual without distractions). While these task-specific differences were much smaller than the global changes from resting state and statistically marginal, the observed pattern matches what would be expected if alpha inflow reduction reflects facilitation. In contrast, power levels in our data did not exhibit a similar double dissociation that would be expected if alpha power were the main mechanism driving attention-related inhibition and facilitation.

One earlier study by Doesburg et al. [67] followed a similar reasoning, trying to assess directed alpha connectivity patterns with transfer entropy to ascertain if they underlie sensory alpha power changes in visual spatial attention tasks. They observed different connectivity changes in a network of regions previously associated with visual spatial attention during the same time window that power modulations take place, but especially a reduction of alpha flow from anterior sites to the contralateral cuneus during attention to a cued hemifield. This closely matches the alpha inflow reduction from the motor/parietal hub we see to the sensory cortices, which increases with attention to the related modality. Wang and colleagues [33], applying Granger causality on EEG measured during a cued visual spatial attention task, equally found a reduction of directed alpha band connectivity from frontal to cue-contralateral occipital brain regions in ‘attend’ vs. ‘ignore’ conditions, though these results were below the statistical threshold required by multiple comparisons correction. Our results however point in the same direction, supporting the interpretation that reductions of alpha outflow from task-relevant higher-order regions to the stimuli-processing sensory cortices reflect an ‘opening’ of relevant processing paths. This is also in line with research on visual hallucinations [70] supporting the interpretation that a decrease of alpha connectivity between higher-order regions and sensory cortices is associated with a disinhibition of the sensory cortex. Moreover, it has been observed that connectivity in the alpha frequency band increases after inhibitory cortical stimulation with low-frequency rTMS [71] and that enhancing alpha connectivity [72] has inhibitory effects on perception [73]. These observations lend further support to our complementary interpretation of alpha connectivity decreases opening functionally relevant processing paths and facilitating processing of information along these paths.

In summary, the dPTE changes in the high alpha band during attention tasks reported here suggest that the gating function canonically ascribed to the alpha band might be realized by mechanisms of connectivity changes in a dynamic network, which could underlie accordant modulations of local alpha power downstream. Specifically, the reduction of flow from higher-order (parietal) or output-related (motor) regions to sensory areas seems to serve a ‘gate-opening’ function or disinhibition, facilitating the transfer and integration of information through the respective pathways.

### Alpha flow from parietal and motor cortex interacts with sensory alpha power in adaptive, goal-oriented network dynamics

We directly assessed the relationship between alpha dPTE and alpha power on a trial-by-trial basis to test for a possible functional relation between them. We found that reduced alpha flow from the motor/parietal hub to primary auditory and primary visual cortex is associated with an alpha power increase within these regions when participants enter in a task-state. This association was absent (in A1) or significantly weaker (in V1) during resting state, pointing to the functional relevance of the relationship for the general task state. Emphasizing the functional relevance of this relationship, the strength of the association between alpha flow from the parietal/motor cortex to the sensory cortices and the modulation of alpha power in the respective sensory cortices was task-specific. In both primary auditory and visual cortex, the strongest relation between reduced alpha inflow and increase of sensory alpha power was found if it corresponded to the task-irrelevant modality (visual task in A1, audio task in V1), and the weakest relation was seen if it was processing the task-relevant modality (audio task in A1, visual task in V1).

In line with the gating-by inhibition literature [2, 74, 75], an up-regulation of alpha power in the task-irrelevant sensory cortex would be expected, and interpreted as inhibition. Interestingly, as mentioned above, averaged power alone does not exhibit here the clear double dissociation of inhibition and facilitation that would be expected by that account. Alpha power averages in both primary sensory cortices increased from resting state with both simple (audio and visual) attention tasks, and for both regions stronger so with the visual task. This might point to a more complex role of sensory alpha power modulations during the distribution of attentional resources than previously thought, which is also suggested by other studies [14, 15, 21, 75], some of which even showed that e.g. the modulation of sensory alpha power can be completely absent dependent on the actual task [18]. In his newest theoretical account, Jensen [15] explores the idea that alpha oscillations are modulated by a secondary mechanism and not under direct top-down control, further discussed below. Alpha power increases are still seen as serving an inhibitory function, but rather emerging as a result of goal-oriented processing, where load and competition of relevant and distractor modalities lead to differing needs and levels of suppression. As a possible communication mechanism for alpha power increases, he suggests, among other options, phasic drives in the alpha band, which have been found to modulate neuronal spiking in intracranial animal recordings [36, 76, 77].

We specifically tested whether alpha dPTE explains or predicts alpha power modulations in sensory cortices, as we were interested in the relationship based on the hypothesis that connectivity patterns or changes of directed in- or outflow could underlie local power modulations (as had e.g. [33, 67], or [15] on a theoretical level). Our linear mixed effects models (LME) assessing the dPTE-power relationship over single trials found the described association of inflow reduction (from a top-down motor/parietal (M/P) hub) and local power increases in the auditory and visual cortices. These models further made the task-specific differences more evident than dPTE or power alone, dissociating clearly between reflecting the task-relevant (attended) and task-irrelevant condition in each sensory cortex. To summarize, while alpha inflow reduction from M/P was slightly stronger in each sensory cortex when its modality was to be attended than when it was to be ignored (and no distraction present), the general inflow reduction from M/P was associated with greater trial-based alpha power increases in the ‘ignore’ condition. From this pattern, it thus cannot be concluded that dPTE drives power in a simple, direct mechanism, in that an inflow reduction is directly reflected in the same level of power increases across conditions. Neither is dPTE flow simply a reflection of power increases in the sender region, as we demonstrated in a final, methodological post-hoc analysis. Rather, the fact that the task- or goal-relevance of a sensory modality underlies both the dPTE changes and even stronger so the dPTE-power relationship (and least visibly so the power averages in our data) suggests a more complex interplay of dynamic mechanisms adjusting information flow, its facilitation and suppression during goal-directed tasks.

### The interplay of goal-dependent global task states and transitory adjustments in a dynamic network

Bringing our observations together, we see a strong case for flow adjustments in the high alpha band (10-13 Hz) as a top-down gating mechanism for the setup of more general, preparatory task states, facilitating and connecting processing paths from relevant sensory input areas and association/selection and output areas - in our case, the primary auditory and visual cortex and the dorsal parietal and motor cortex of the output-relevant left hemisphere. The main mechanism of this setup, here, is a strong reduction of the alpha outflow from the association and motor output regions that is constant and high at resting state, apparently serving a ‘gate opening’ function or facilitation. In addition, the same connections show small, but meaningful adjustments in connection strength related to task-demands, i.e. the modality to be attended and reacted to with a motor response shows further inflow reduction, suggesting further facilitation or prioritization of its input. At the same time, during the tasks, a relationship forms between the reducing alpha flow from the higher-order regions to the sensory cortices and increasing alpha power in those sensory cortices. This relationship, then, also exhibits meaningful adjustments in strength across the different task conditions, such that when modality suppression is task-relevant, the trial-based dPTE-associated power increases are stronger.

An interesting case regarding the fine-grained adjustments of both M/P to sensory alpha dPTE flow and dPTE-power association is the visual task with auditory distractions. As we did not have a clear hypothesis for this condition, the following discussion is more speculative. In this task, alpha power levels in both sensory cortices have the same mean as during resting state, while they increase for the other motor-response tasks. However, the trial-based dPTE-power association shows an intermediate strength in each sensory cortex, while the association is weaker when the cortex modality is task-relevant and stronger when task-irrelevant (and without distractions in both these cases). This would indicate that the fluctuation of both auditory and visual alpha power levels over trials in this task is, in contrast to resting state, happening in association with equally transitory dPTE adjustments of an overall intermediate strength between each facilitated/simple and suppressed/simple condition, respectively. At the same time, both simple audio and visual tasks without distraction showed greater overall reductions of M/P-to-sensory alpha flow in both sensory cortices than the visual task with auditory distraction. Taken together, these results do not seem random and could reflect some kind of bottom-up competition of the incoming stimulation from both modalities. As the auditory distractions in this task were more numerous than the visual modulations to be detected, they possibly interfered with the visual facilitation (less reduction of alpha flow to visual cortex; more visual alpha power increase with inflow reduction), all the while being more inhibited than during the easier and more straightforward visual task (less inflow reduction to auditory cortex and stronger auditory alpha power increase with inflow reduction). As stated above, this interpretation is speculative, as we had no clear informed hypothesis about this particular control condition. The results here are suggestive, however, and it could be worthwhile in future experiments to test alpha connectivity changes with more specifically designed distractions in both auditory and visual modalities systematically manipulating levels of load or difficulty for comparison.

While we did not systematically introduce or investigate load in the present study, the concept is increasingly regaining relevance in the research on neural resource allocation [75, 78]. Jensen [15] makes a case for load and biased competition as the relevant concepts underlying the goal-oriented processing adjustments, which modulate alpha oscillations and result in inhibitory alpha power increases in turn. Connecting our findings to that account, we can support that modulations of alpha oscillations reflect goal-oriented processing. By looking at oscillatory changes both in alpha connectivity and alpha power, we see on the one hand a set of stronger and more global changes in alpha connectivity, possibly related to a general, goal-oriented task state or preparatory network setup that distinguishes our attention tasks from resting state, opening channels for relevant sensory input and attentional processing and integration, as well as motor output (if task-relevant). On the other hand, we see more small-scaled and possibly more transitory or dynamic adjustments in alpha connectivity along the same connections and alpha power modulations in the sensory cortices happening in accordance with those, both of which seem to have functional relevance related to the facilitation or prioritization of task-relevant vs. inhibition of task-irrelevant or competing information, respectively.

The size and patterns of these more fine-grained modulations of alpha oscillations, reflected in both connectivity and associated power, could also be related to load and competition. The generally low load of the stimulation in our purposefully basic tasks, for example, might be one reason the canonically expected power dissociation did not emerge clearly. This same low load, together with the almost identical stimulation in the audio and visual task (and modulation and response windows cut out) could be why the task-specific differences in the dPTE reductions are meaningful in tendency, but small in scale. That our pool of trials consists of time segments cut out between the modulations (and button responses) in an ongoing stimulation - and with varying distances in time to these - might further predispose us to see more global, preparatory task-related changes than short-term, stimulus-related ones. In contrast to the typical visual spatial attention paradigms, which use short trials where attention is cued and switches often between hemifields, our participants just had a simple and global attention instruction (target modality) for each sub-task, consisting of four 100s-long trials with modulations (prompting a button-press) occurring at random intervals within these. Competition with a more pronounced distractor load is present in our study only in the visual task with auditory distractions as discussed above. While the concept seems meaningful for explaining the observed pattern, we cannot draw further conclusions here, as it was not assessed in a systematic way.

Returning to the question of how neural resource allocation and information flow might be managed in the brain, we think that a) oscillations in the alpha band play a crucial role, and b) different functional mechanisms in the alpha band dynamically work together to optimize processing resources and information flow for a specific goal or task. Some of these seem to happen on a more global task-oriented scale, setting up (and maintaining) a preparatory network state. We identified here reduced alpha flow from higher-order association or output regions to relevant input regions as one major mechanism for opening and prioritizing processing paths and their integration. Clearly, further studies need to show how universal this mechanism is, or how specific to the kind and details of our attention task set and the regions it recruits. Other mechanisms seem to reflect more short-term adjustments in the global preparatory network state, possibly reflecting modulatory updates over time and trials in the network setup for short-term goal changes, and through feedback mechanisms for ongoing optimization in neural recruitment and information flow for task goals based on preceding trials (e.g. small-scale updates in attentional focus, response strategy, adjustments based on changing load etc.). We believe that the functional smaller-scale differences we see between our attention sub-tasks in both alpha inflow reductions and related local alpha power increases in the sensory areas based on their changing task-relevance reflect such dynamic adjustments in the global task state.

Future studies may elucidate further how alpha oscillations help goal-oriented, prioritized processing via both changes in alpha flow between regions and local alpha power increases within circumscribed regions, how these neural phenomena enact cognitive mechanisms acting both on a more global, preparatory scale as well as on a more short-term, performance-adjusting scale, and how these mechanisms interplay. Studying changes in alpha connectivity in addition to power measures seems to be worthwhile, as does a more systematic focus on how differences in task designs shape the goal-oriented recruitment and interaction of mechanisms for long-term preparation and short-term adjustment.

## Conclusion

Moving from resting state to a set of tasks requiring attention in different modalities and related responses produces significant changes in directed whole-brain connectivity, with major qualitative changes in the high alpha band (10-13Hz) appearing as central for task-specific functional adjustments. We posit here a global-scale mechanism of prioritizing and integrating task-relevant processing paths between relevant sensory input regions (auditory and visual cortex), higher-order association (dorsal parietal cortex), and output regions (motor cortex) in the left hemisphere, with motor connections only affected in the tasks requiring motor responses. Specifically, task-related global reductions of the strong alpha outflow from the parietal and motor to the sensory cortices present at rest may serve the function of ‘opening’ gates to facilitate information flow and create a preparatory network state for the tasks. Between sub-tasks, smaller-scale modulations in the global reductions of motor/parietal-to-sensory flow seem to further facilitate the attended modality, and correlate with stronger alpha power increases in the unattended modality. We suggest that the task- and region-specific reduction of connectivity in the alpha band reflects a disinhibition of relevant communication channels serving the optimization of goal-oriented information flow in the brain, acting on both a more global, general scale for the tasks, as well as serving more short-term task-specific adjustments, where it interacts with related adjustments of local levels of alpha power.

## Acknowledgements

We would like to thank Martin Kaltenhäuser, Carolin Spielau-Romer, and Antonia Keck for assistance in data collection. This research was funded in entirety by the Deutsche Forschungsgemeinschaft (DFG MU 3916/1-1 Emmy-Noether-Programm). We declare no conflicts of interest.

---

**Figure S1:**
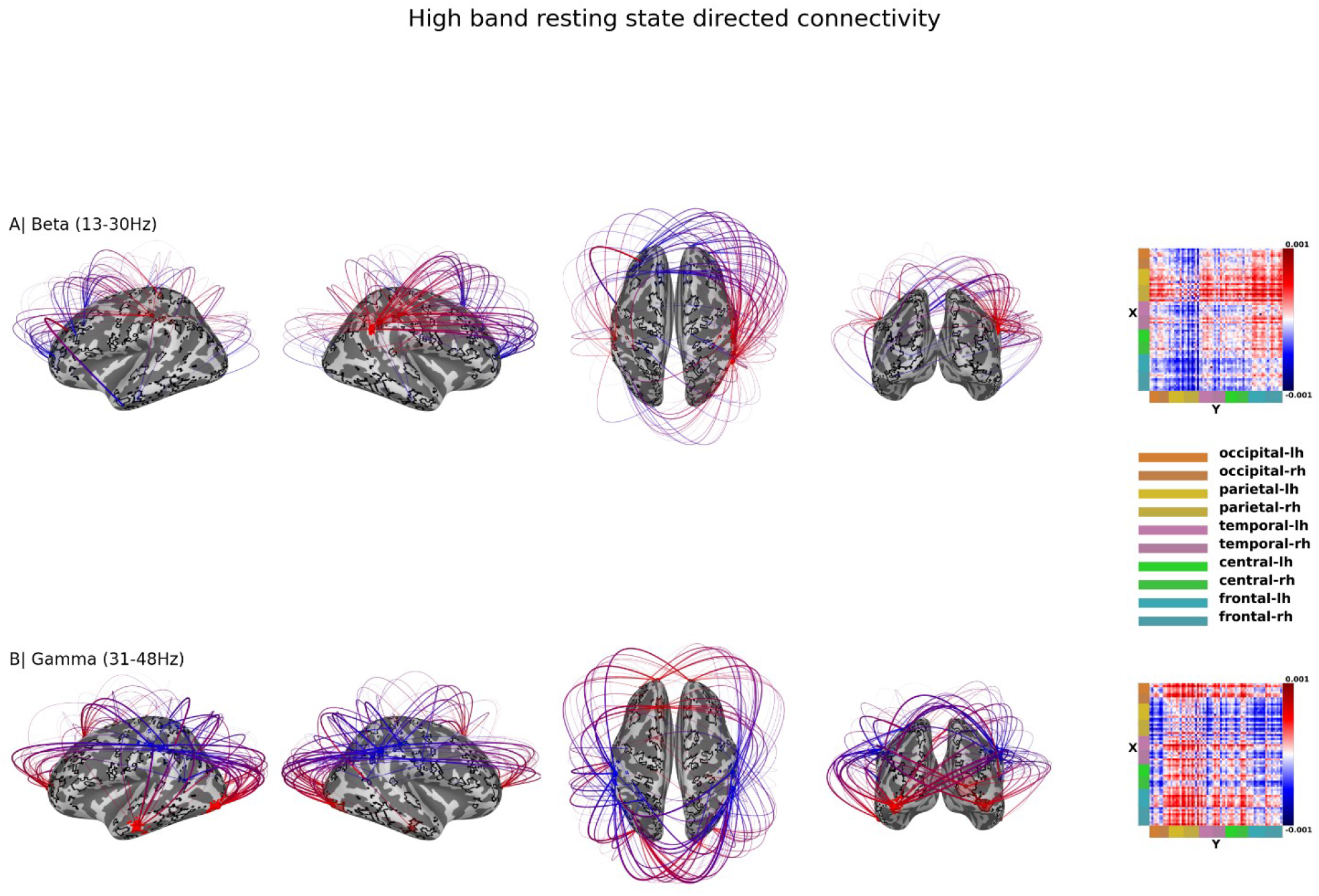
The top 150 strongest connections. Information movement from source to destination is indicated by red to blue. Matrix values represent dPTE values for resting state, subtracted from 0.5 to centre them around 0. A) Resting state beta band (13-30Hz) connectivity B) Resting state gamma band (31-48Hz) connectivity.

